# Competitive pre-ordering during planning persists in kinematically fused sequential movements

**DOI:** 10.1101/2025.08.18.670892

**Authors:** Helena Wright-Wieckowski, Jason Friedman, Joseph M. Galea, Katja Kornysheva

## Abstract

Results in human and non-human primates have shown that elements of a movement sequence are pre-ordered in parallel competitively before execution, a process known as competitive queueing (CQ). However, it is unclear whether the preplanning of individual movements persists in continuous skilled actions that involve greater biomechanical integration and is associated with the formation of new motor primitives (neural fusion). We investigated how kinematics impact sequence planning in a handwriting-like task asking whether fusing velocity curves between adjacent movements affects movement preparation. Participants were trained and tested for two days to perform two sequences of four sequential centre-out-and-back movements from memory in a delayed sequence production task using a stylus on a Wacom tablet. To manipulate kinematic fusion between subsequent strokes, participants were assigned to one of three groups that were trained to perform the sequences either with acute, right or obtuse angles between sequential targets. Probe trials assessed the availability of constituent movement elements for fast and accurate execution towards each target during planning. Movement elements associated with later sequence positions were less available than earlier movements, regardless of kinematic fusion, in line with CQ findings for discrete typing sequences in humans. Importantly, a more pronounced CQ gradient was associated with higher fusion, faster initiation and greater accuracy of sequence production. These findings indicate that kinematically fused sequential actions do not result in the formation of new movement primitives (neural fusion) with a single movement plan. Instead, they continue to be planned separately and are associated with skilled performance.

**New & Noteworthy:** Using a handwriting-like sequencing task, we manipulated kinematic fusion through target geometry and examined the relative availability of individual movements towards targets as a marker of competitive queuing (CQ) during planning. Contrary to the assumption that behavioural fusion reflects their neural fusion of movements into a new motor primitive, we show that even under high levels of fusion, the underlying sequence elements remain competitively pre-ordered by their sequence position benefitting performance.

## Introduction

Executing a fast and accurate series of movements from memory is essential to everyday skilled actions in humans, in particular dextrous movements such as typing, handwriting and drawing. How does the brain control naturalistic sequential movements that have become accurate and fluent following training?

Neurophysiological findings in non-human primates have demonstrated that when drawing geometrical shapes after long-term training, the neural representation of each stroke segment in the prefrontal cortex is retrieved simultaneously during planning rather than serially (Averbeck et al., 2002). Similarly, learning two-element sequential reaches led to the primary motor cortex preparing the second reach even before the first reach was completed, further supporting the concept of parallel planning (Zimnik and Churchland, 2021).These findings have been replicated in humans through discrete typing tasks using MEG pattern decoding and behavioural probe tasks, with both techniques providing evidence for parallel planning of upcoming movements in sequences produced from memory (Kornysheva et al., 2019; Mantziara et al., 2021). The strength of the pattern was shown to reflect the preplanning of the upcoming sequence and readiness to perform the sequence (Mantziara et al., 2021).

Computational modelling shows that the parallel planning of sequential movements occurs through a competitive process (Houghton, 1990; Houghton and Hartley, 1996; Burgess and Hitch, 2005). Multiple items are active simultaneously in the competitive planning layer, with the most active one being selected for output. Once selected, each item is temporarily inhibited, allowing the next most active item to be chosen, after which it gradually recovers from inhibition. This computational model, known as competitive queuing (CQ), effectively explains order errors in serial recall (Burgess and Hitch, 1992; Burgess and Hitch, 2005). Additionally, the CQ model accounts for learning new sequences by proposing that different sequences recruit different context signals. This mechanism also accounts for the Hebb repetition effect, as repeated exposure to a sequence strengthens its representation, making it easier to recall sequence orders over time (Burgess and Hitch, 2006).

While previous research in non-human primates and humans has primarily focused on discrete movement sequences, such as typing or straight-line reaches performed at acute angles, well-trained dextrous movement sequences often combine discrete elements with smooth continuous movements, like in handwriting, drawing and tool use. Whilst discrete movement sequences exhibit a velocity profile with distinct deceleration phases, resulting in brief pauses between movements, continuous movements co-articulate or kinematically fuse two movements into a single velocity peak to reduce jerk and energy expenditure (Todorov and Jordan, 1998; Kornysheva, 2016; Friedman and Korman, 2019; Zacks and Friedman, 2020; Sporn et al., 2022; Sporn and Galea, 2025). The 2/3 power law suggests that the velocity of execution of a handwriting/drawing movement is directly related to the curvature of the trajectory, with velocity increasing as the radius of curvature increases (Lacquaniti et al., 1983). This directly relates to the minimum jerk model, which suggests that the motor system minimises mean squared jerk (changes in acceleration) – to optimise smooth movement (Flash and Hogan, 1985). Fusion occurs as a result of reducing the mean squared jerk, which is optimised at different training stages and is a sign of skilled performance (Friedman et al., 2013; Sporn et al., 2022). In a series of studies, Sosnik et al. (Sosnik et al., 2004, 2007, 2015) examined this idea by having participants perform a handwriting-like task, moving through a sequence of points with varying obtuse and acute angles. Fusion occurred at the obtuse angles, as reflected in the velocity profiles, which shifted from four distinct peaks to two as individual movements gradually merged. This process can be facilitated by prolonged practise (Sosnik et al., 2015), reward (Sporn et al., 2022), as well as through movement observation (Friedman and Korman, 2019) and the use of analogies (Zacks and Friedman, 2020).

Beyond differences in physical trajectories, studies suggest that the control mechanisms underlying continuous, fused movements may differ from those of discrete movements (Zelaznik et al., 2002; Spencer and Ivry, 2005; Kornysheva, 2016). This links to the idea that as kinematic fusion increases, the motor system progresses from the control of individual sequence element to the formation of a new holistically planned movement (Sosnik et al., 2015; Zimnik and Churchland, 2021). Accordingly, if kinematic fusion is accompanied by neural fusion, the individual movements should combine into a single movement primitive, with distinct planning patterns corresponding to this new fused action. Discrete strokes toward sequential targets would no longer feature in the movement plan. In contrast, if kinematic fusion merely reflects the temporal overlap or co-articulations of separate movements, the constituent movements would remain distinct and discrete strokes to targets pre-planned competitively according to their position in the sequence (CQ).

Here, we investigated the influence of increased kinematic fusion in a handwriting-like sequence task on sequence planning, focusing on the CQ of upcoming sequence elements. We adapted the complex learning task from Sporn et al. (2022), training participants over two days to produce 8 centre-out sequences from memory. By varying target angles, where more obtuse angles promote greater fusion, we aimed to assess whether kinematic fusion alters movement primitives, with consequences to the planning of elements that become part of a new kinematic movement element. If new movement primitives emerged, the acute-angle group, with the lowest fusion, would show the most pronounced CQ gradient, as individual elements remain distinct. Conversely, the obtuse-angle group, with the highest fusion, would show a disrupted CQ gradient, reflecting merged elements that could no longer be independently probed. Instead, we found that even highly fused movements were planned as separate elements, with CQ associated with faster and more accurate performance.

## Materials and Methods

### Participants

Fifty-four participants took part in two sessions across two days and were randomly assigned to one of three experimental groups: acute-angle (N=19, M=22.3 years, SD=5.85), right-angle (N=16, M=18.9 years, SD=5.73), obtuse-angle (N=19, M=20.5 years, SD=6.22). The number of participants for each condition was based on the group sizes used in a previous study investigating movement planning (Mantziara et al., 2021). Eighteen additional participants were excluded due to not completing all two sessions (N=2), a technical error (N=1), or a performance error rate above 50% in Probe trials or an error rate exceeding three standard deviations above the mean for Sequence trials during either the test phase or the post-test (N=16). Handedness was assessed using the Edinburgh Handedness Inventory (Oldfield, 1971) and all participants were right-handed according to the laterality quotient (acute: M=88.2, SD=15.3; right: M=86.9, SD=15; obtuse: M=80.8, SD=18) and had normal or corrected-to-normal vision, with no neurological, developmental or psychiatric conditions and no injury to the hand or wrist. None of the participants had previously participated in a similar sequence production experiment. For financial reimbursement, participants received £10 for their time or for non-financial reimbursement, participants received 3 SONA credits. Participants were recruited via the research participation scheme (RPS) and the Centre for Human Brain Health (CHBH) mailing list. The study was approved by the University of Birmingham ethics committee (ERN_09-528P) and all participants provided informed consent before participating.

### Apparatus

Participants were seated comfortably in a quiet booth on a static chair in front of a desk with a Wacom One graphics tablet display (screen size: 33.6cm x 22.2 cm, sampling rate: 240Hz). To better simulate the experience of handwriting, participants were permitted to adjust the angle and move the display on the desk in front of them until they found a comfortable and natural handwriting-like position. It was adjusted at the start of the experiment until participants could reach all targets with the stylus pen comfortably from the centre-point by flexing and extending the fingers whilst the wrist was kept on the same spot of the display and without any visual obstruction by the hand. Participants were encouraged to support their hand during handwriting by touching the screen with the ulnar side of the hand, as they would naturally do on a piece of paper.

They received initial verbal instructions, where they were told the sequences to perform and what to expect for each block, before performing 2D reaching movements with a stylus using their right hand, which closely resembled everyday handwriting movements, e.g. like handwriting the letter “K”, but involving different angles and a different number of strokes. The visual stimuli were presented on the display, and the stylus tip position on the display surface was sampled using the Repeated Measures Toolbox created by (Friedman, 2014) (https://github.com/JasonFriedman/RepeatedMeasures/tree/v1.0.0). The experiment was run using Psychtoolbox.

### Experimental Design

We implemented a sequence preparation and production paradigm involving handwriting-like movements using a pen stylus on a tablet based on the delayed sequential production paradigm involving finger presses (Mantziara et al., 2021). Participants were trained to produce two sequences of four centre-out and back movements with a stylus from a central target and performed from memory as fast and accurately as possible (Figure 1a). Both the central and outer targets had a diameter of 0.4 cm, with the centre point of the four outer targets placed from the centre point of the central target at a distance of 5cm. Participants were <u>assigned to different groups that were</u> trained to perform the sequences either with acute, right-angle or obtuse angles between sequential targets. The acute-angle group (45°) targets were positioned at angles of 57.5°, 77.5°, 102.5°, and 122.5° relative to the horizonal line through the centre target. The right-angle group (90°) included targets positioned at 35°, 55°, 125°, and 145°. Finally, the obtuse-angle group (140°) had targets positioned at 5°, 35°, 145°, and 175°. The angle of each condition (45°, 90° and 140°) reflected the angle between targets 1 & 3 and 2 & 4 (Figure 1b).

**Figure 1.**
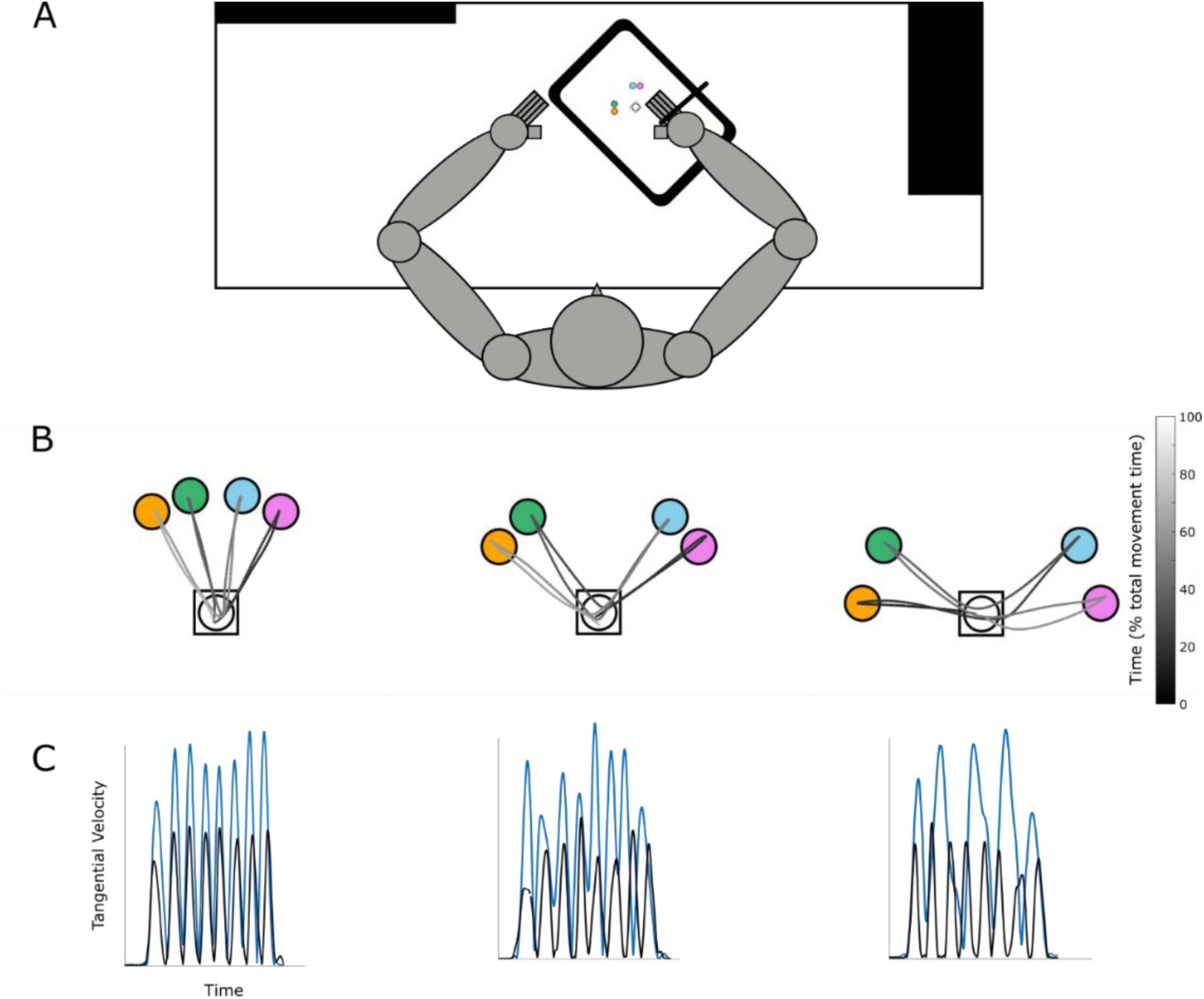
Experimental set-up. **A:** Participants used a stylus to perform centre-out handwriting movements on a pen display, with hand and display positioned individually for comfort. Their wrist and hand rested on the screen, mimicking natural handwriting on paper. **B:** An example of the trajectories for each condition (acute, right-angle, obtuse), with the greyscale bar representing time relative to the overall movement time (% movement duration). **C:** Trial examples of tangential velocity profiles normalised by overall movement time (% movement duration) for each condition during the early training phase (black) and the late test phase (blue) scaled by movement duration.

In the Sequence trials, participants were first instructed to place their stylus on the centre target. After 300ms, the sequence cue (either A or B) appeared for a fixed duration of 400ms. Following a delay of 1000ms, the Go cue appeared around the centre target, signalling participants to perform the sequence from memory. For Probe trials, the trial began in the same way as Sequence trials but instead of the Go cue, the centre cue changed colour to match one of the target colours. Participants were then prompted to make a single movement from the centre target to the outer target of the corresponding colour as quickly and accurately as possible (Figure 2).

**Figure 2.**
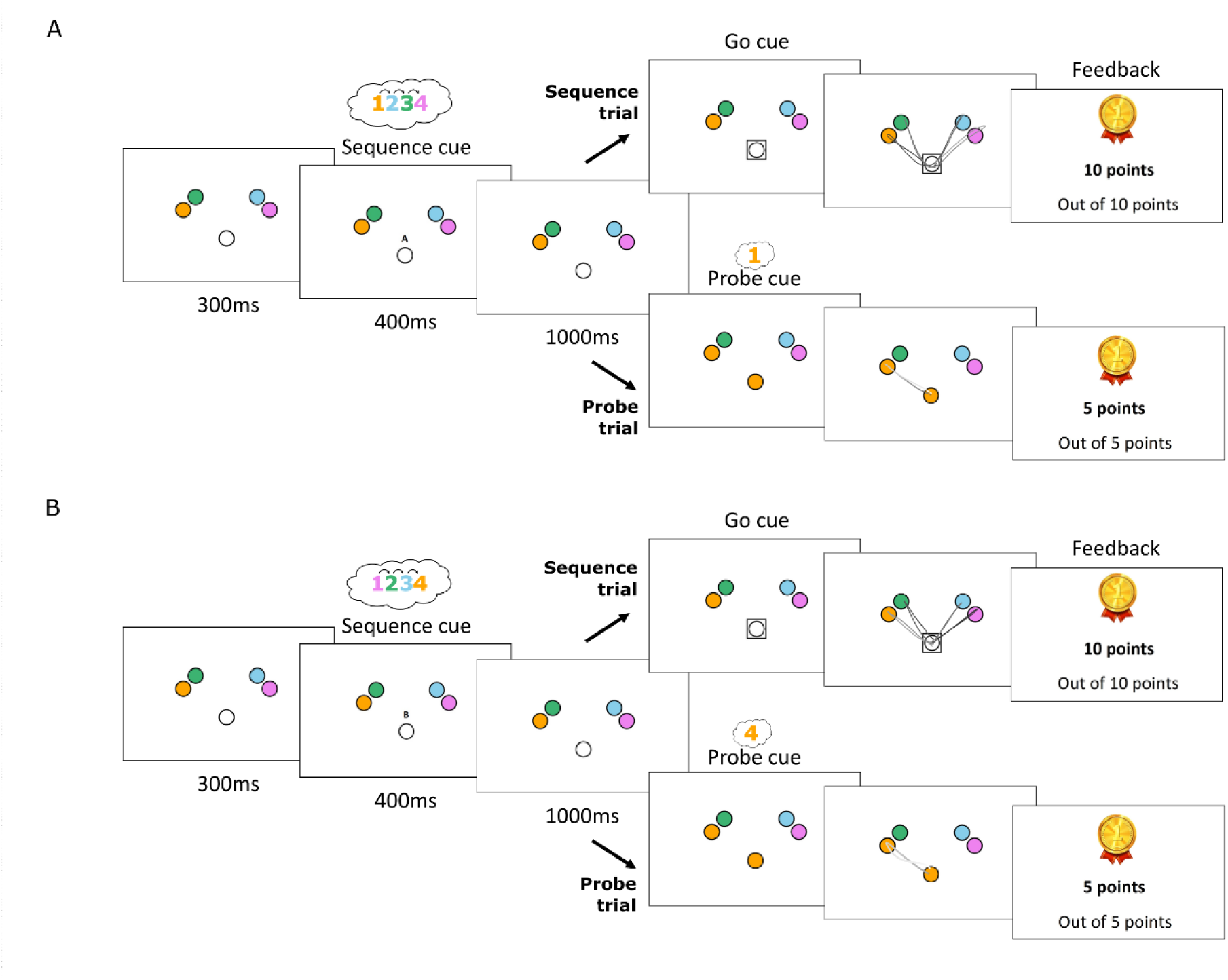
Diagram of each sequence and trial type. **A:** An example of a Sequence and Probe trial. The sequence begins when the stylus is positioned in the centre target, signalling the start of the trial. After a delay of 330ms, the sequence cue appears above the centre target and remains visible for a fixed duration of 1000ms before the go cue is presented. Once the go cue appears, participants produce the sequences from memory for Sequence trials and perform a single centre-out movement for Probe trials. Feedback is then provided based on how quickly they initiate the sequence/single movement following the go/probe cue. **B:** An example of sequence B, the mirror of sequence A.

Once they returned to the centre, they received feedback. Points were awarded based on the speed and accuracy with which participants initiated the sequence, with rewards contingent on completing the sequence in the correct order. On Sequence trials, participants could earn up to 10 points, while on Probe trials, a maximum of 5 points was available (Figure 2). To determine the reward, each correct trial’s reaction time (RT) was compared to the RTs from the preceding 20 trials of the same type (sequence or probe). The RTs were ranked, and the current trial’s percentile determined the number of points awarded, for example, RTs within the fastest three received the maximum reward. After each trial, the screen went blank for 500ms, during which participants lifted their pen off the screen. Participants received feedback based on their RT after each trial. At the start of the pre-test block, participants had not yet completed 20 Sequence trials, so for the initial ranking, all 20 comparison times were set to 5000ms as a baseline. If a trial was not completed or the trial timed out (after 20 seconds) before the participant performed the correct order, then the participant would receive 0 points. The trial continued until they completed the sequence correctly or until it timed out, after 20 seconds for Sequence trials, and after 1 second for Probe trials. The timeout for Probe trials was shorter due to the shorter movement involved and to encourage rapid responses.

The experiment was conducted over two days, with the first day dedicated to training participants on the sequences, and the second day focused on testing their performance (Figure 3). This two-day structure allowed participants to significantly increase fusion of movement strokes (Sporn et al., 2022; Sporn and Galea, 2025). Due to a technical error, 11 participants in the right-angle group and 14 in the obtuse-angle group had an unequal number of Probe trials across sequence positions, during the test phase. To address this, we equalised the data by identifying the lowest number of trials across positions and excluding any additional trials beyond this number in the other positions for each participant.

**Figure 3.**
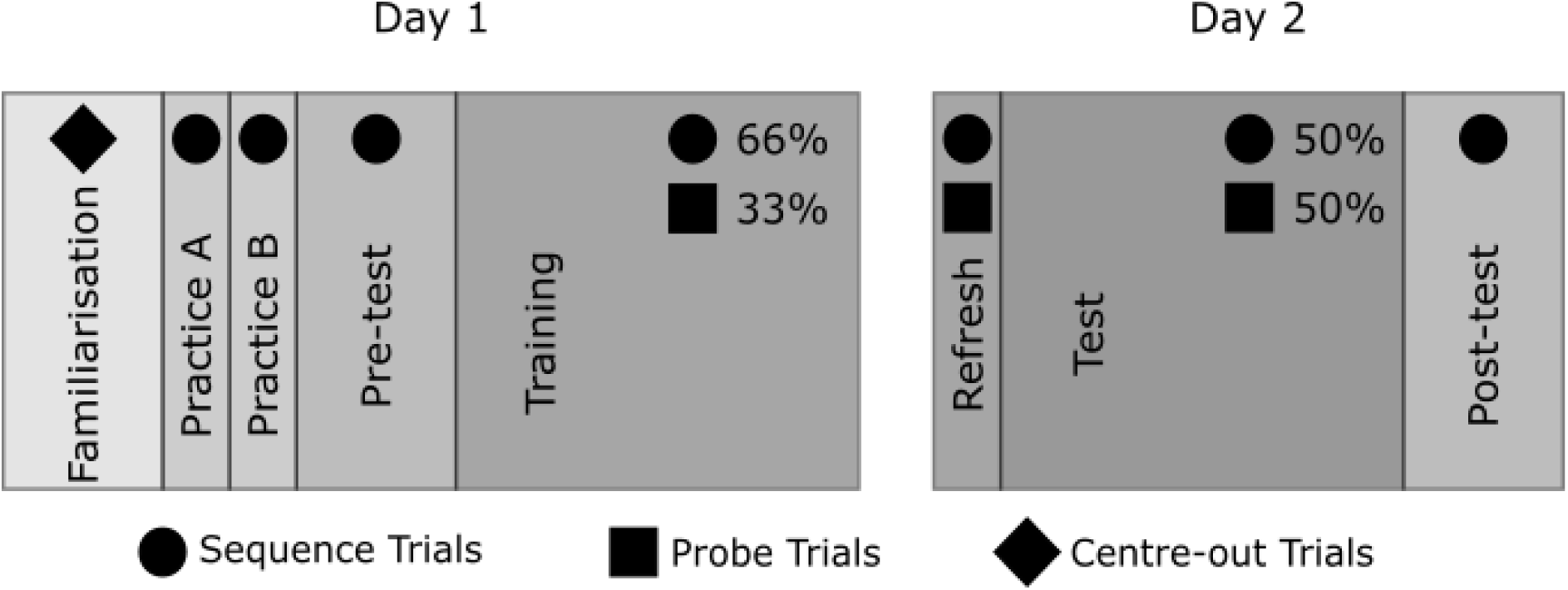
Schematic showing the type of trial for each block across the two-day study.

### Procedure

Day 1 consisted of several phases - familiarisation, practice, sequence pre-test and training. To familiarise the participants with the tablet and moving the stylus in the task space, the familiarisation block included 20 trials of single centre out-and-back movements to each target. Trials were randomly distributed for each target. Next, to familiarise participants with the production of the two sequence orders, they were shown written instructions on the target order for both sequence A and sequence B. During the two subsequent practice blocks participants were instructed to produce sequences A and B in a blocked manner. Each block contained 8 trials of either sequence A or B, respectively, with blocks counterbalanced across participants. All participants were instructed to complete each trial as quickly and accurately as possible. If participants completed the sequence in the wrong order, they would have to complete the sequence straight after without restarting the full trial until the order was correct or until the trial timed out after 20 seconds. Trials where sequences were not produced correctly after the delay period were regarded as erroneous trials. This was the same for all Sequence trials in the experiment. The subsequent sequence pre-test was designed to enhance the familiarity with both sequences and as a comparison to the post-block to evaluate sequence learning. Participants completed 20 Sequence trials, with 10 trials for each sequence, presented in a randomized order. In the subsequent training phase, participants completed 15 blocks of 24 trials each: 16 Sequence trials (2 sequences x 8 repetitions) and 8 Probe trials (2 sequences x 4 positions) for day 1, i.e. 240 (66.7%) Sequence trials and 120 (33.3%) Probe trials. After each trial, participants received points based on their performance. Probe trials were introduced during the training blocks so that participants would become familiar with them.

Day 2 followed a similar design as in Mantziara et al (2021). First, participants completed a refresher block consisting of 24 trials of 66.7% sequence, 33.3% probe. This was followed by a test phase which included 14 blocks of 16 test trials, consisting of 8 (50%) sequence production trials (2 sequences x 4 repetitions) and 8 (50%) Probe trials (2 sequences x 4 positions) per block, amounting to 112 sequence production trials and 28 Probe trials for each sequence position across the two sequences. The post-test was the same as the pre-test. It was included initially because we aimed to obtain a measure of fusion which would be compared to the pre-test block. However, as participants were still learning the sequences in the pre-test, they had a very low accuracy rate on average (M=37.27%, SD=27.37). Additionally, pre- and post-tests only contained Sequence trials, which led to faster RTs compared to blocks that also included Probe trials like the Test phase. To address this, we used the first training block as the baseline measure of performance and the end of the test phase as an equivalent comparison.

### Data analysis

Data analyses were performed using MATLAB (Version: 9.15.0 (R2024a)) and RStudio (Posit team (2025). RStudio: Integrated Development Environment for R; Posit Software, PBC, Boston, MA. URL http://www.posit.co/.). The WACOM stylus tip position data was filtered using a 4^th^ order, two-way low pass Butterworth filter, with a cutoff of 10 Hz, prior to following data analyses.

#### Sequence production

The Sequence trials produced from memory from the test phase were used to analyse sequence production. Specifically, we examined the fusion index (Fusion), calculated by comparing the maximum tangential velocities of two adjacent peaks *V_max1_* and *V_max2_* to the minimum velocity between them *V_min_* and normalised by the average peak velocity between the two peaks (Equation 1).

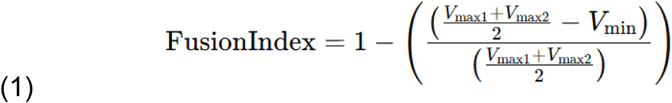

Smaller differences indicate greater movement fusion, i.e. more continuous sequence execution (Sporn et al., 2022). Fusion was calculated for correctly executed trials only and for all apart from the first and last two velocity peaks in the sequential movement as these belonged to centre to target out-and-back movements of the first and last targets in the sequence which a priori cannot be fused. A mean was calculated across each trial.

Furthermore, sequence initiation time (InitRT) was defined from the Go cue to the point at which the participant reached 5% of their maximum tangential velocity during the increase in velocity to the first velocity peak, marking sequence movement onset. Movement time (MT) was measured from this point until the last time where the tangential velocity exceeded 5% of its maximum and occurred within the final half-second of the movement as defined by the stylus touching the screen; when both conditions were met, it signified the end of the movement. Accuracy was calculated as the percentage of correctly executed trials in blocks of interest. Dwell time was the amount of time each participant spent in each target and was calculated by determining the coordinates of points along the target’s circumference, based on its centre coordinates and radius. We then identified the times at which the participant first crossed the circumference (entry) and subsequently exited it (exit) and computed the dwell time by subtracting the exit time from the entry time. For each Sequence trial, the total dwell time across all targets was calculated and averaged.

The absolute values to quantify learning were taken from the first block of the training phase and the last 2 blocks of the test phase (Figure 4). The initial training block contained 16 Sequence trials and 8 Probe trials, but in the test phase, there were 8 Sequence trials and 8 Probe trials. To compensate for the disproportionate number of trials, we included the last two blocks of the test phase as a comparison to the first training block.

**Figure 4.**
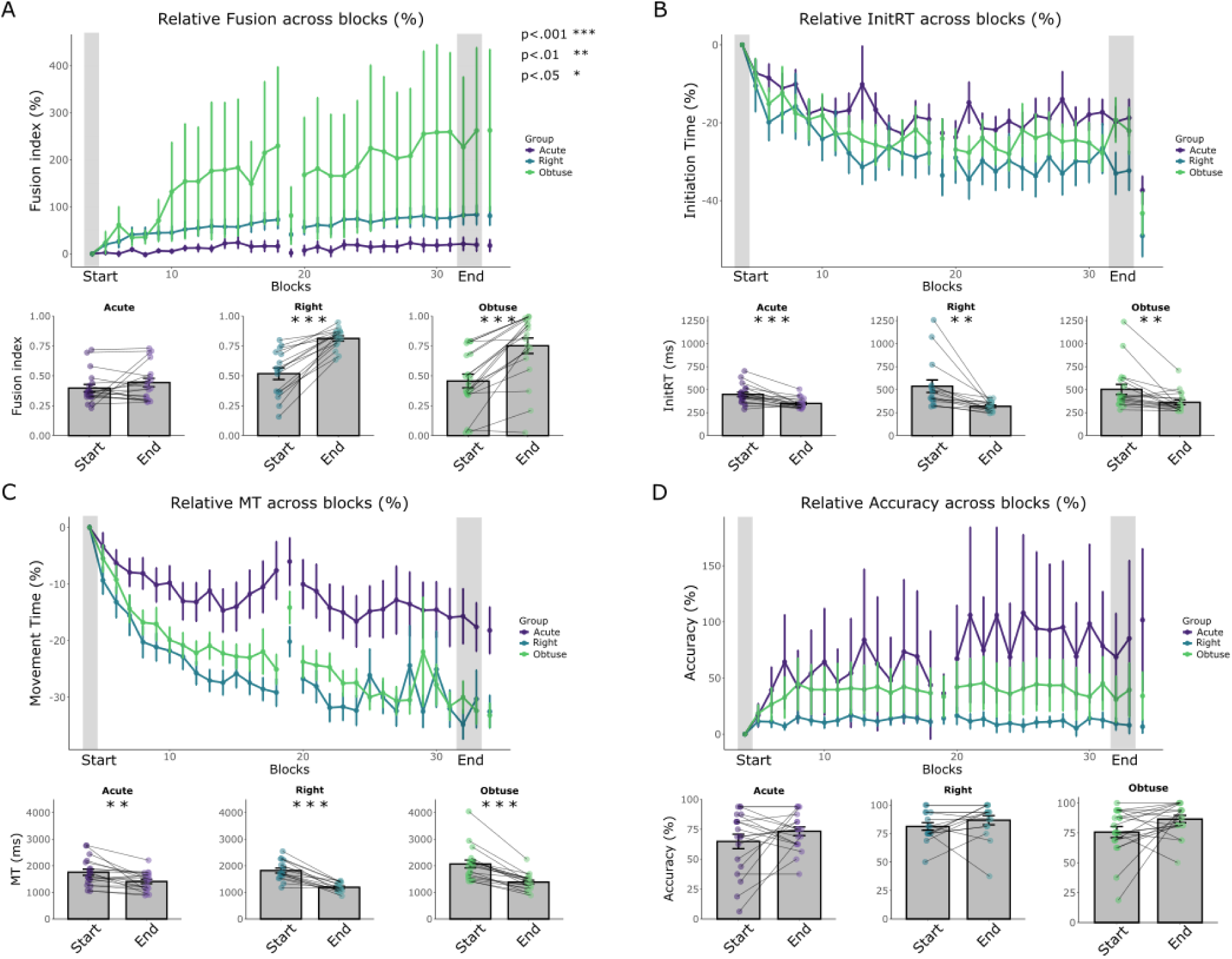
Learning performance metrics across groups. **A:** Change of fusion score in percent for correct Sequence trials compared to the first training block across all blocks for each group (acute, right-angle, obtuse), upper panel. The first training block (grey rectangle) contains 16 trials, while the final two test blocks on day 2 (grey rectangles) contain 8 trials each. Error bars represent the standard error of the mean. Lower panel, comparison of RT for each group between the first training block (start) and the final two test blocks (end). **B:** Change of InitRT for correct Sequence trials across all blocks for each group, upper panel. Comparison of RT for each group between the start and end block, lower panel. **C:** Change of MT for correct Sequence trials. **D:** Change of accuracy percentage.

#### Sequence planning

In Probe trials, participants were prompted to move to a specific target as quickly and accurately as possible, following a sequence cue and a preparation delay. The target was indicated by the central target changing to a colour that matched the colour of one of the outer targets (Fig. 2). To measure the availability of the upcoming movements in the sequence for fast and correct execution, we first calculated the median reaction time (RT) across correctly executed Probe trials (ProbeRT), as well as the error rate across Probe trials (ProbeError). Since each Probe trial contained 2 movements (to the target and back), and a trial was classified as an error if it contained more than 2 peaks, indicating the participant initially started moving in a different direction before correcting the movement, or if the RT was negative i.e. movement started before the Go cue. The ProbeRT was measured from the onset of the Probe cue to when the participant entered the target which was equivalent to the RT measure used in the button press tasks (Mantziara et al., 2021).

To test the interaction between the factors of position (1^st^, 2^nd^ 3^rd^, 4^th^) and group (acute, right, obtuse angle) in Probe trials, raw RTs and error rates were analysed using a mixed-model ANOVA. Significant interaction effects were followed up with a one-way ANOVA within each position to examine group differences, and paired t-tests within each group to examine position differences. In addition, sequential contrasts (2–1, 3–2, 4–3) were used to assess consecutive position differences within each group, and these differences were then compared across groups to evaluate interaction effects. For each participant, we calculated the percentage increase in RT and error rate for each probed position relative to the first position. This enabled us to visualise the position-dependent increases relative to the first position across conditions (Figure 6a). Next, we calculated the mean relative differences in RT and error rate between adjacent positions (i.e. the fourth minus the third position, the third minus the second position and the second minus the first position) for each participant. This served as an index of movement pre-ordering during sequence planning (ProbeRT, ProbeError; Figure 6b). Slower reaction times and higher percentage of errors associated with later positions would indicate CQ, suggesting that each movement was competitively pre-planned.

Furthermore, a median split was calculated based on each performance measure (sequence InitRT, MT, fusion index, accuracy, dwell time) for raw median ProbeRTs and mean ProbeError rates for each position.

To further investigate the relationships between sequence performance (sequence InitRT, MT, fusion index, accuracy, dwell time), planning (ProbeRT, ProbeError), and relative performance (i.e., the percentage increase from the first training block to the end of the test phase), we conducted a permutation-based correlation analysis using Spearman’s rho with 5000 iterations (Figure 8a).

#### Inferential statistics

To test the data for normality, we used the Shapiro-Wilk test and then the Levene’s test to measure the homogeneity of variance. Additionally, for the probe data analysis, we tested the assumption of sphericity using Mauchly’s test. When sphericity was violated, Greenhouse-Geisser corrections were applied.

The fusion index, InitRT, MT and accuracy were analysed using a two-way mixed ANOVA design to examine the difference between group (acute, right, obtuse angle) and blocks (Start: defined as the start of training, End: defined as the end of the test phase). Significant interaction effects were investigated using a one-way ANOVA for each block to assess group differences. P-values were adjusted using the Holm correction, and paired t-tests were used to evaluate differences between blocks within each group. We used the measures for fusion index, InitRT, MT and accuracy from the test phase to examine sequence production (Figure 5). We did a one-way ANOVA to compare the means of these variables across groups. For post-hoc tests, we checked for pairwise differences between groups using Tukey HSD. A stepwise linear regression was performed to examine which performance variables were predictors of ProbeRT and ProbeError. Predictors were removed if they did not improve the model fit (Figure 7c).

**Figure 5.**
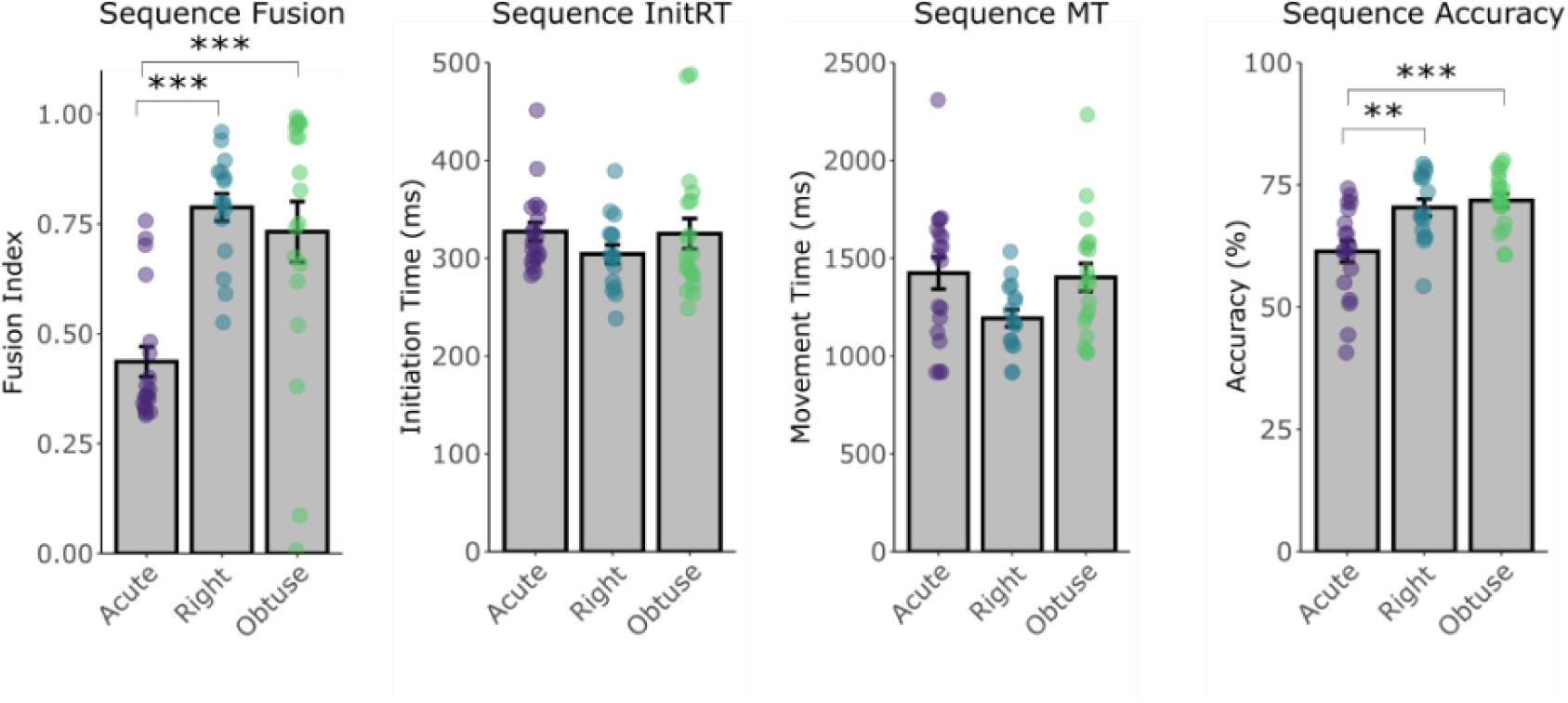
Absolute sequence production performance across the test phase. The mean fusion index, InitRT (ms), MT (ms), and accuracy (%) for correct Sequence trials during the test phase of the experiment on day 2 for each group.

**Figure 6.**
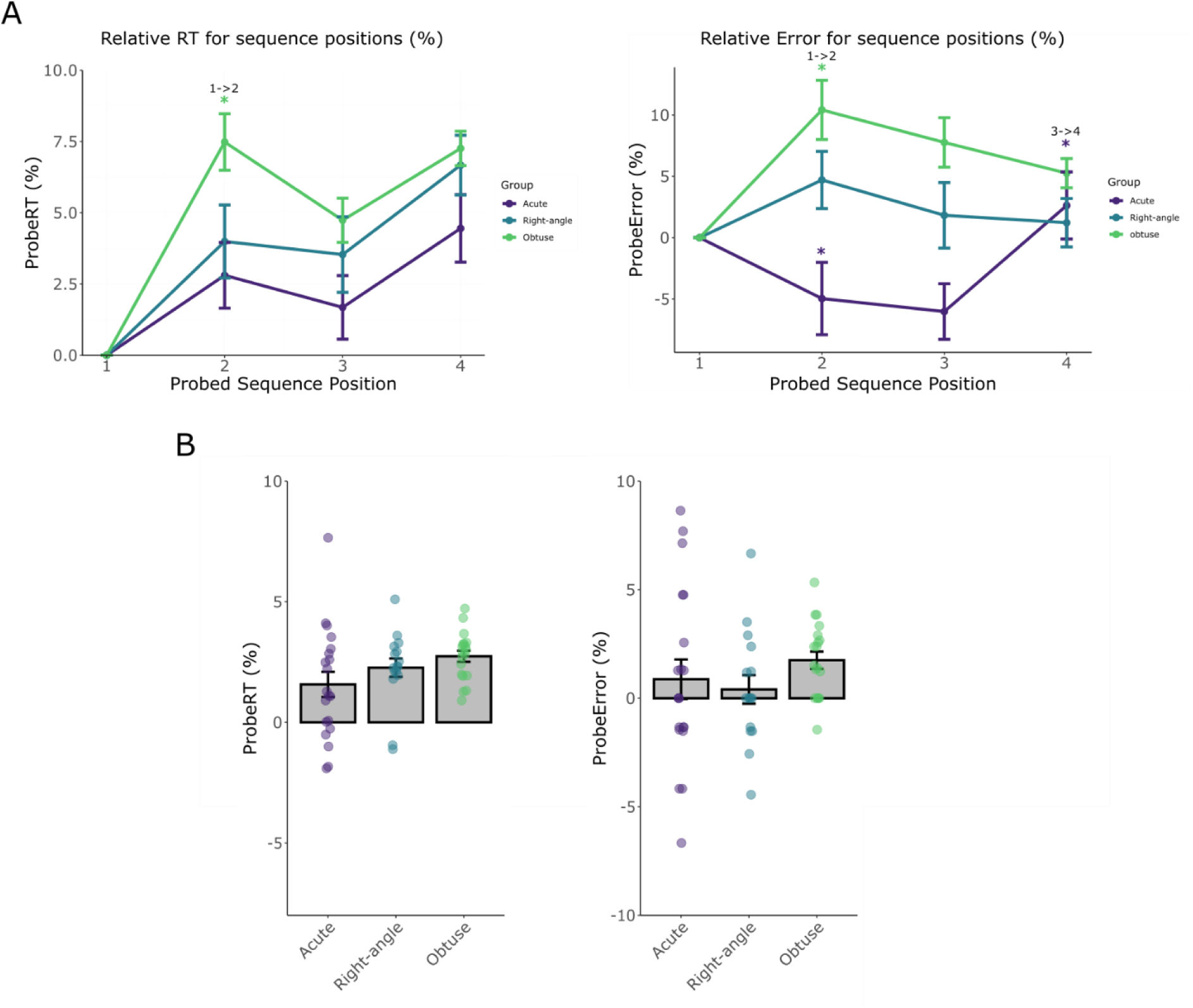
Competitive queuing during planning split by group. **A:** The relative RT and error of Probe trials to the first position was measured for each sequence position for all groups. **B:** The CQ scores for ProbeRT and ProbeError.

**Figure 7.**
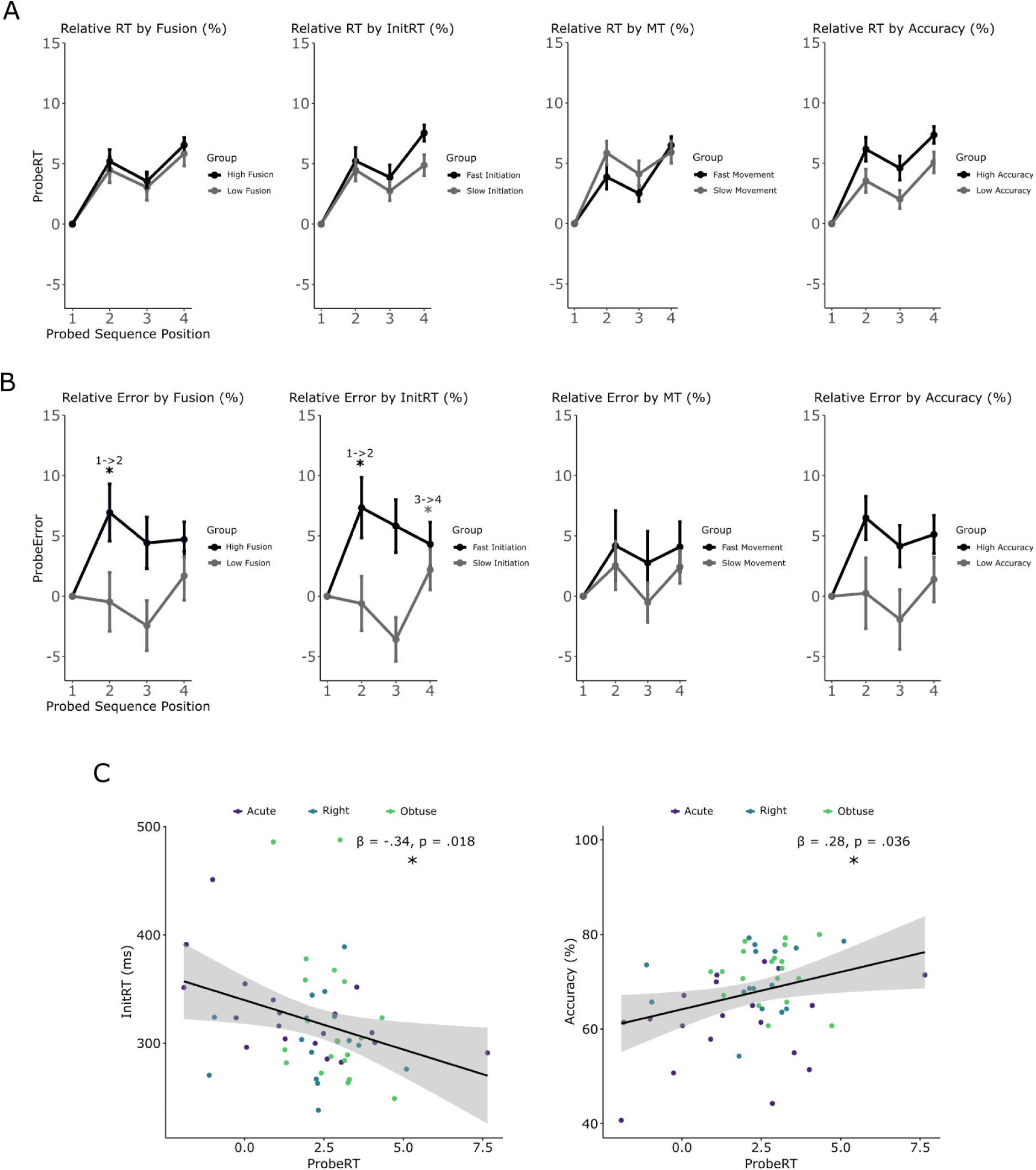
Pre-ordering of upcoming movements during planning split by performance. **A:** The relative reaction time of Probe trials to the first sequence position of correct trials, grouped by fusion, initiation, MT and accuracy, during the test phase following a median split. **B:** The relative errors for each sequence position. **C:** Scatter plots showing the relationship between InitRT and accuracy with ProbeRT, based on the predictors identified through stepwise linear regression.

**Figure 8.**
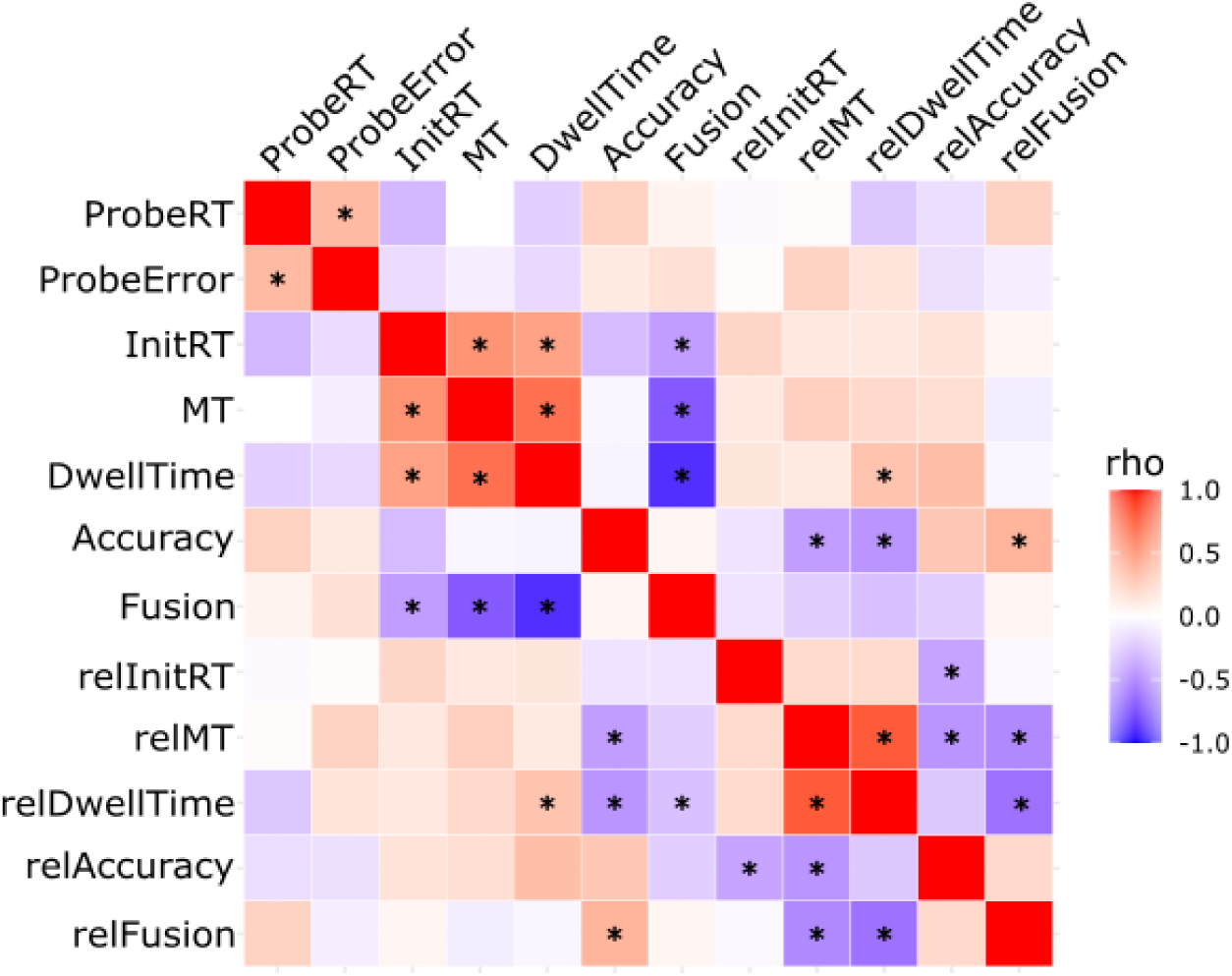
Correlations between sequence planning, production and training related changes in performance. Correlation matrix showing sequence planning (ProbeRT, ProbeError) and the Fusion scores, InitRT (ms), Dwell time (ms), MT (ms), Accuracy (%) and the relative values. These measures were taken from the test phase and averaged across the corresponding blocks. Significant correlations after permutation-based FDR correction (p<.05) are marked with *.

To examine the reduction in variability of fusion scores within each block as an additional marker of learning, we examined the standard deviation of fusion (Figure 9a). Similarly to learning, we conducted a two-way mixed ANOVA design to examine the difference between group and blocks (Figure 9b) and then grouped by fusion (Figure 9c).

**Figure 9.**
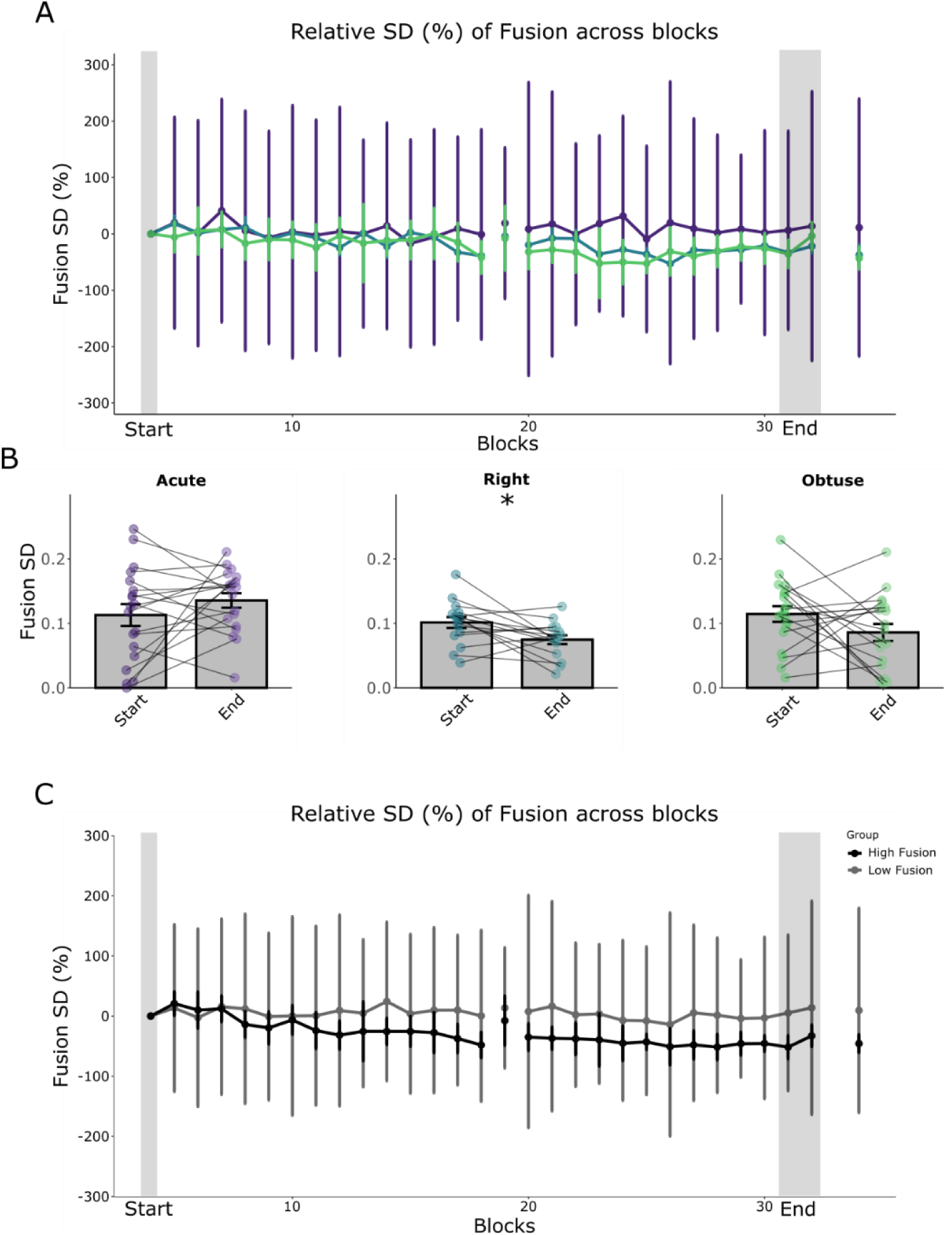
Variability in fusion scores for each group. **A:** Relative change in standard deviation of fusion scores to the first training block across all blocks for each group (acute, right, obtuse angle). The first training block (grey rectangle), the last two test blocks (grey rectangle). Error bars represent standard error. **B:** Comparison of standard deviation for each group between the first training block (start) and the final two test blocks (end). **C:** Relative change in standard deviation scores, grouped by fusion following a median split.

## Results

### Right-angle and obtuse-angle groups increased movement fusion with training

First, we investigated the changes in performance from the start of training to the end of the test phase across the two-day experiment. We found no difference in fusion scores between groups at the start of training, *F*(1, 51) = 9.06, *p* < .001, η²ₚ = .26, acute (*M* = .40, *SD* = .14), right-angle (*M* = .52, *SD* = .19), and obtuse (*M* = .46, *SD* = .75). We found an increase in fusion from the first block (*M* = .45, *SD* = .20) to the last block (*M* = .66, *SD* = .25), *F*(1, 51) = 87.02, *p* < .001, η²ₚ = .63. This was dependent on the experimental group, *F*(2, 51) = 13.67, *p* < .001, η²ₚ = .35. The interaction was driven by the acute angle group showing no significant increases in fusion (*M* = .40, *SD* = .14 to *M* = .44, *SD* = .15; *p* = .29, *d* = –.30), whereas the right-angle (*M* = .52, *SD* = .19 to *M* = .81, *SD* = .087; *p* < .001, *d* = – 2.08) and obtuse group (*M* = .46, *SD* = .25 to *M* = .75, *SD* = .28; *p* < .001, *d* = –1.51) showed increased fusion from start (start of training) to end (end of test phase). There was no significant difference in fusion between right and obtuse angle groups at the end of training (*p* = .37, *d* = .29). Thus, as hypothesized, the larger angles in the right and obtuse angle groups in the handwriting sequences, enabled the participants to kinematically fuse subsequent movements as they became more skilled at sequence production.

### Sequence initiation time, movement speed and accuracy reduced independently of target angle group

We found a general decrease in InitRT across groups, *F*(1, 51) = 34.82, *p* < .001, η²ₚ = .41, *M* = 494.07 ms, *SD* = 213.2 ms in the first block to *M* = 345.05 ms, *SD* = 80.4 ms in the last block, with no main effect of group, *F*(2, 51) = 0.35, *p* = .71, η²ₚ = .014, or an interaction with group, *F*(2, 51) = 1.78, *p* = .181, η²ₚ = .065. This was because, as expected, all groups became faster in initiating the handwriting-like sequence after the Go cue. Similarly to InitRT, we found a general decrease in MT across groups, *F*(1, 51) = 97.27, *p* < .001, η²ₚ = 0.66, from *M* = 1887 ms, *SD* = 532 ms to *M* = 1338 ms, *SD* = 306 ms, with no main effect of group, *F*(2, 51) = 1.59, *p* = .21, η²ₚ = 0.059, or an interaction with group, *F*(2, 51) = 3.57, *p* = .035, η²ₚ = 0.12. We found an increase in accuracy from the first block, *F*(1, 51) = 6.67, *p* = .013, η²ₚ = 0.12, *M* = 73.50%, *SD* = 21.8% to the last block, *M* = 81.94%, *SD* = 16.1%, with no main effect of group, *F*(2, 51) = 5.717, *p* = .006, η²ₚ = 0.18, and no interaction with group, *F*(2, 51) = 0.23, *p* = .80, η²ₚ = 0.009.

### Absolute fusion and accuracy were more pronounced in the right- and obtuse-angle groups compared to the acute-angle group

To assess absolute performance, we calculated fusion, InitRT, MT, and accuracy over the full test period, which was also used for analysing Probe trial performance. There was a difference between groups with regard to absolute fusion scores across the whole test period, *F*(2, 51) = 14.67, *p* < .001, η²ₚ = .37, with the acute group having a smaller fusion index (M = .437, SD = .15) compared to the right (M = .788, SD = .12; *p* < .001, *d* = –2.54) and obtuse-angle group (M = .732, SD = .30; *p* < .001, *d* = –1.25), but no difference between the right-angle and obtuse group (*p* = .72, *d* = .23). We found no differences for absolute InitRT, *F*(2, 51) = 1.030, *p* = .36, η²ₚ = .039, or MT, *F*(2, 51) = 3.14, *p* = .052, η²ₚ = .11. However, groups differed in sequence accuracy, *F*(2, 51) = 10.27, *p* < .001, η²ₚ = .29, with the acute group (M = 61.35%, SD = 9.49%) having a lower accuracy compared to the right- (M = 70.36%, SD = 7.06%; *p* = .003, *d* = –1.06) and the obtuse-angle groups (M = 71.80%, SD = 5.80%; *p* < .001, *d* = –1.32). There was no difference in accuracy between the right and obtuse angle group (*p* = .84, *d* = –.23). In summary, as hypothesized, target angles between adjacent movements resulted in increased absolute fusion scores for right- and obtuse-angle groups. However, accuracy was lower for the acute-angle group, likely reflecting the greater precision needed due to the close proximity of targets.

### Planning handwriting-like sequences involves competitive queuing of individual trajectories

Next, we examined whether the competitive queuing of upcoming movements during planning can be observed behaviourally in a handwriting-like task, whereby the movement availability of sequence element (CQ) would be modulated by their sequence position, as observed in discrete movement sequences (Averbeck et al., 2002; Kornysheva et al., 2019; Mantziara et al., 2021). Importantly, we set out to test whether CQ is altered by the degree of kinematic fusion. Specifically, we expected kinematic fusion to lead to neural fusion of movements adjacent in the sequence. The centre-out movements that featured in the original sequence and assessed in Probe trials would stop being part of the sequence leading to a reduction or even a breakdown of CQ. Accordingly, this would be more pronounced in the right- and obtuse-angle group where kinematic fusion could occur.

RT in trials that probed the availability of centre-out movements corresponding to different parts of the sequence was modulated by sequence position, *F*(3, 153) = 37.75, *p* < .001, η²ₚ = .425, and interacted with experimental group, *F*(6, 153) = 2.38, *p* = .032, η²ₚ = .085.This interaction was mainly driven by an increase from position 1 to position 2, the obtuse group had a significant increase compared to the right angle group, t(153) = 2.39, p=.036, and the acute group, t(153) = 3.42, p = .0031. There was no difference between the acute and right-angle group, t(153) = .80, p=.42. There were no significant differences between groups for any of the positions when the *p*-values were adjusted using the Holm correction: position 2 (*p* = .01, η²ₚ=.13), position 3 (*p* = .10, η²ₚ=.13), and position 4 (*p* = .10, η²ₚ=.13). The absolute RT values were modulated by group, *F*(2, 51) = 3.50, *p* = .038, η²ₚ = .121. When examining the absolute RT results within groups, we found no increase from position 1 to 2 for the acute group (*M* = 471.81 ms, *SD* = 40.33 ms to *M* = 485.34 ms, *SD* = 43.89 ms; *p* = .095, d=-.59). However, there was an increase for the right-angle group (*M* = 499.14 ms, *SD* = 32.36 ms to *M* = 518.60 ms, *SD* = 36.82 ms; *p* = .029, d=-.77) and the obtuse-angle group (*M* = 490.00 ms, *SD* = 50.26 ms to *M* = 527.14 ms, *SD* = 61.96 ms; *p* < .001, d=-1.73). From position 2 to 3, there was no difference observed for the acute (*M* = 485.34 ms, *SD* = 43.89 ms to *M* = 479.27 ms, *SD* = 43.13 ms; *p* = .35, d=.36) and right-angle groups (*M* = 518.60 ms, *SD* = 36.82 ms to *M* = 516.23 ms, *SD* = 35.56 ms; *p* = .71, d=.095), although there was a decrease for the obtuse-angle group (*M* = 527.14 ms, *SD* = 61.96 ms to *M* = 513.15 ms, *SD* = 54.45 ms; *p* = .015, d=.73). From position 3 to 4, we found no difference for the acute (*M* = 479.27 ms, *SD* = 43.13 ms to *M* = 493.47 ms, *SD* = 38.79 ms; *p* = .13, d=-.53) or obtuse-angle groups (*M* = 513.15 ms, *SD* = 54.45 ms to *M* = 525.54 ms, *SD* = 54.76 ms; *p* = .052, d=-.56); however, there was an increase for the right-angle group (*M* = 516.23 ms, *SD* = 35.56 ms to *M* = 532.06 ms, *SD* = 35.29 ms; *p* = .029, d=-.81). From position 1 to 3, there was no difference for the acute (*M* = 471.81 ms, *SD* = 40.33 ms to *M* = 479.27 ms, *SD* = 43.13 ms; *p* = .35, d=-.31) or right-angle groups (*M* = 499.14 ms, *SD* = 32.36 ms to *M* = 516.23 ms, *SD* = 35.56 ms; *p* = .040, d=-.65), although a significant increase was found for the obtuse-angle group (*M* = 490.00 ms, *SD* = 50.26 ms to *M* = 513.15 ms, *SD* = 54.45 ms; *p* < .001, d=-1.38).

We found that there was an increase in error as the position increased, *F*(2.48, 126.35) = 4.42, *p* = .009, η²ₚ = .080, which was dependent on the experimental group, *F*(2, 51) = .35, *p* = .71, η²ₚ = .013. The interaction effect was mainly driven by a difference between position 1 and position 2, the acute group had a decrease compared to the right-angle group, t(153) = 2.88, p = .009, and the obtuse angle group, t(153) = 5.01, p<.001. The obtuse group had a larger increase compared to the right-angle group, which was marginally significant, t(153) = 1.9, p =.058. The acute group had a larger increase from position 3 to position 4 compared to the right-angle group, t(153) = −3.11, p=.0045, and the obtuse group, t(153) = −3.91, p<.001. There was a difference between groups for position 1 (*p* = .0025, η²ₚ = .27) and position 4 (*p* = .024, η²ₚ = .23). For position 1, *F*(2, 51) = 8.53, *p* < .001, there was a difference between the acute (M = 9.43%, SD = 7.50%) and right-angle group (M = 4.81%, SD = 4.65%; *p* = .017), and between the acute and obtuse group (M = 2.16%, SD = 3.32%; *p* < .001). For position 4, *F*(2, 51) = 5.38, *p* = .024, there was a difference between the acute (M = 12.81%, SD = 7.57%) and the right-angle (M = 1.50%, SD = 1.32%; *p* = .002), and between the acute and obtuse angle group (M = 1.89%, SD = 1.59%; *p* = .011). The error values were not modulated by group, *F*(2, 51) = 0.89, *p* = .41, η² = .03416.

When examining within-group differences between positions, we found for position 1 to 2 there was a difference for the obtuse group (M = 0.579%, SD = 0.90% to M = 3.21%, SD = 2.82%; *p* = .003, *d* = –0.96), but there was no difference for the acute (M = 2.63%, SD = 2.09% to M = 1.63%, SD = 2.11%; *p* = .55, *d* = 0.32) or right-angle group (M = 1.19%, SD = 1.17% to M = 2.38%, SD = 1.89%; *p* = .19, *d* = –0.56). For position 2 and 3 there was no difference for any of the groups: obtuse (M = 3.21%, SD = 2.82% to M = 2.53%, SD = 2.39%; *p* = .37, *d* = 0.32), right (M = 2.38%, SD = 1.89% to M = 1.69%, SD = 1.82%; *p* = .17, *d* = 0.60), or acute angle groups (M = 1.63%, SD = 2.11% to M = 1.26%, SD = 1.28%; *p* = .55, *d* = 0.29). For position 3 to 4 there was a difference for the acute group (M = 1.26%, SD = 1.28% to M = 3.58%, SD = 2.12%; *p* = .001, *d* = –1.07), but not for the right (M = 1.69%, SD = 1.82% to M = 1.50%, SD = 1.32%; *p* = 1.00, *d* = 0.13) or the obtuse angle group (M = 2.53%, SD = 2.39% to M = 1.89%, SD = 1.59%; *p* = .37, *d* = 0.28).

Overall, this suggests that the obtuse group had a more pronounced CQ gradient. When we calculated the CQ gradient as a score for RT and error (Figure 6b), we found no significant differences for RT, *F*(2, 51) = 1.80, *p* = .18, η²ₚ = .066, or error, *F*(2, 51) = 0.87, *p* = .43, η²ₚ = .033.

In sum, we found support for the CQ of upcoming movements during planning of handwriting-like sequence elements. In contrast, we did not find support for kinematic fusion resulting in a reorganisation of sequence planning through CQ. In contrast right and obtuse angle groups that showed more kinematic fusion merging some of the elements in the sequence also had a more pronounced CQ of the original centre-out elements in the sequence.

### Better performance was associated with more pronounced pre-ordering of upcoming movements to targets

Despite significant group effects related to target angle, fusion and other performance measures varied considerably within each group. For example, some participants in the obtuse-angle group showed no fusion even after two days. To assess whether CQ was related to performance, we conducted a median split based on individual performance measures rather than on group membership: fusion (low: *M* = 0.416, *SD* = 0.16; high: *M* = 0.86, *SD* = 0.098), InitRT (slow: *M* = 354.6 ms, *SD* = 50 ms; fast: *M* = 284.46 ms, *SD* = 18.2 ms), MT (slow: *M* = 1575.48 ms, *SD* = 249 ms; fast: *M* = 1119.21 ms, *SD* = 150 ms), and accuracy (high: *M* = 74.78%, *SD* = 3.15%; low: *M* = 61.12, *SD* = 7.23).

When the groups were split by fusion index, there was no interaction between fusion and positional increase for ProbeRT scores, *F*(3, 156) = 0.086, *p* = .97, η²ₚ = .002 but there was a significant interaction for ProbeError scores, *F*(2.19, 114.03) = 3.16, *p* = .042, η²ₚ = .057. For ProbeError this was driven by a larger increase from position 1 to position 2 for those who had greater fusion, t(156) = 2.78, p=.008. Overall, this suggests that participants who tend to fuse more also show a more pronounced preordering of movements for first versus later movements in the sequence.

When the group was split by sequence InitRT, the positional increase was not modulated for ProbeRT, *F*(3, 156) = 1.10, *p* = .35, η²ₚ = .021, but only for ProbeError, *F*(2.22, 115.19) = 5.91, *p* = .003, η²ₚ = .10. The latter interaction was again driven by those with faster InitRTs having a larger increase from position 1 to position 2, t(153) = −2.94, p=.004, whereas those with slower InitRTs did not show any CQ from the first movement and only had an increase from position 3 to position 4, t(153) = 2.73, p=.007. These findings suggest that overall participants who were faster at initiating sequences also had a more pronounced preordering of the first versus the later elements in the sequence.

A more pronounced preordering of movements was not linked to individuals having slower and faster MT during sequence execution. There was only a marginal trend in ProbeRT, F(3, 156) = 2.17, p = .09, η²ₚ = .040, and ProbeError showed no relationship with MT, F(2.1, 109.42) = 0.62, p = .55, η²ₚ = .012.

When the groups were split based on sequence accuracy, there was only a marginally significant interaction between ProbeRT and accuracy, *F*(3, 156) = 2.21, *p* = .089, η²ₚ = .041, and no significant interaction for ProbeError and accuracy, *F*(2.14, 111.29) = 1.90, *p* = .15, η²ₚ = .035.

To test for a linear relationship between performance variables and the amount of CQ, a stepwise linear regression was conducted to examine whether the individual performance scores (Fusion, InitRT, MT and Accuracy) predicted the ProbeRT and ProbeError scores. InitRT (β = −.34, *p* = .018) and accuracy (β = .28, *p* = .036) were retained as significant predictors of ProbeRT. The final model was statistically significant, *F*(1, 58) = 4.11, *p* = .011, and explained 15% of the variance in RT (*R*² = .198, adjusted *R*² = .150). For ProbeError, the final model retained no predictors, only the intercept differed significantly from zero (*B* = 1.09, *SE* = 0.41, *t*(53) = 2.64, *p* = .011). This suggests a linear relationship between performance variables related to sequence initiation speed and accuracy and the order-specific pre-ordering of upcoming movements during planning.

### Fusion correlates with markers of skilled performance

In an exploratory follow-up analysis, we examined the relationship between planning variables (ProbeRT and ProbeError), performance measures (InitRT, MT, Dwell time, Accuracy, and Fusion), using relative measures as proxies for learning-related changes in performance (relative change in InitRT, MT, Dwell time, Accuracy, and Fusion from start of training to the end of the test phase). This analysis employed Spearman’s rank-order correlations, with significance levels estimated via a permutation test (5,000 iterations).

ProbeRT was positively correlated with ProbeError, *r*ₛ(56) = .33, *p* = .045, confirming that relative availability to quickly and correctly produce the probed movements are markers of a common underlying competitive queuing (CQ) mechanism, although not identical. No correlations between ProbeRT or ProbeError and performance or learning variables reached the permutation threshold.

Although many performance variables were interrelated, we specifically highlight correlations with kinematic fusion, since the latter was the focus of the experimental manipulations in this study. Specifically, fusion correlated negatively with InitRT (*r*ₛ(52) = – 0.44, *p* < .006), MT (*r*ₛ(52) = –0.72, *p* < .001) and Dwell time (*r*ₛ(52) = –0.84, *p* < .001), showing that greater fusion reflects faster initiation and a faster and smoother execution of the handwriting-like sequence. Increase in fusion from start of training (relFusion) was positively correlated with Accuracy, *r*ₛ(52) = 0.40, *p* = .012 and negatively correlated with relMT (*r*ₛ(52) = –0.70, *p* < .001) and relDwell (*r*ₛ(52) = –0.69, *p* < .001). This suggests that training-related changes went hand-in-hand with higher sequence execution accuracy and improvements in speed and fluency.

Overall, fusion emerged as a key indicator of skilled performance. While the strength of the association between CQ scores, fusion and other performance measures linked to higher skill varied across CQ markers, all significant effects consistently indicated that greater pre-ordering of movements was advantageous for skilled performance.

### Right- and obtuse-angle groups show a reduction in fusion variability with training

We wanted to evaluate the consistency of movement trajectories and to understand whether coarticulation occurred from the outset or whether fusion developed and stabilised with practice. To investigate this, we examined the variability in fusion scores over time by measuring the standard deviation of fusion. We found a main effect of group, *F*(2, 51) = 4.18, *p* = .021, η²ₚ = .14. There was no main effect of block, *F*(1, 51) = 1.21, *p* = .28, η²ₚ = .023, and there was a marginally significant interaction between group and block, *F*(2, 55) = 3.01, *p* = .058, η²ₚ = .11. This weak interaction effect was mainly driven by a decrease from early training to the test phase for the right-angle group (*M* = .10, *SD* = .034 to *M* = .075, *SD* = .027; *p* = .02, *d* = .65). There was no decrease for the acute (*M* = .11, *SD* = .074 to *M* = .14, *SD* = .049; *p* = .23, *d* = −.28) or for the obtuse angle group (*M* = .11, *SD* = .053 to *M* = .09, *SD* = .057; *p* = .15, *d* = .34), likely because of the variability in fusion in the obtuse-angle group, with two participants not showing any fusion. We therefore ran a post hoc analysis based on the median split in fusion and found a main effect of group, *F*(1, 51) = 12.43, *p* < .001, η²ₚ = .20, and no main effect of block, *F*(1, 51) = 1.09, *p* = .30, η²ₚ = .021. There was an interaction between group and block, *F*(1, 51) = 10.19, *p* = .002, η²ₚ = .17. This was mainly driven by a significant decrease in fusion SD for high fusers (*M* = .11, *SD* = .049 to *M* = .070, *SD* = .039; *p* = .004, *d* = .59); there was no decrease found for low fusers (*M* = .11, *SD* = .061 to *M* = .13, *SD* = .047; *p* = .16, *d* = –.29). This suggests that fusion is a learning-related effect, and it was not pre-existing coarticulation that was occurring from the outset.

## Discussion

Skilled daily activities rely on fast, accurate and smooth sequences of movements, exemplified by tasks such as handwriting, drawing, music and sports performance. The goal of our study was to examine how kinematic fusion of movements adjacent to each other in a sequence affects the planning of the underlying sequence structure. Here, we show that upcoming movements to targets are competitively queued (CQ) during motor planning in both discrete and continuous handwriting-like movements. Contrary to the assumption that behavioural fusion is a marker of neural fusion, we find that behavioural fusion does not reduce CQ and tends to be more pronounced for participants showing more skilled performance.

More fusion between targets adjacent in the sequence did not result in a disrupted CQ of these elements during planning. Thus, continuous movement trajectories between targets without distinct dips in velocity may not be taken to reflect neural fusion or a fused sequence planning strategy. Our planning data suggests that the underlying structural elements of a sequence (centre-out-centre movements) remained intact and echoed those of single movements assessed in Probe trials, despite kinematic differences between movements as part of continuous sequential and discrete single trajectories. On the contrary, participants who performed the sequences with greater kinematic fusion also tended to show a stronger modulation of the sequence elements by their relative position, i.e. a more pronounced CQ.

While the preservation of individual components within kinematically fused movements is surprising for skilled continuous movements, findings from animal electrophysiology suggest that markers of neural fusion of extensively practiced compound movements remain elusive, particularly in humans and non-human primates. When monkeys produced multiple highly practised two-reach sequences of movements, muscular activity formed one continuous pattern, however, the neural representation in the primary motor and dorsal premotor cortex did not show evidence of fusion into a dedicated holistic representation, with neural patterns fully overlapping with those for single reaches or reaches with a long gap between them (Zimnik and Churchland, 2021). Further, the second reach was planned simultaneously with the execution of the first reach. This suggests that in the cortical motor system, producing rapid sequences does not draw on movement fusion and is in line with previous research on parallel competitive movement planning of discrete movements prior to their execution (Averbeck et al., 2002; Kornysheva et al., 2019; Mantziara et al., 2021). In another study with macaques performing a double-reaching task, it was found that when both sequential reaches were planned together during preparation (Wang et al., 2024), one signal was modulating the other, so that a multiplicative model better captured neural activity in the motor cortex. This suggests that the planning of the second sequential reach is not just unfolding alongside the first but is actively modulating the second reach. While a temporary multiplicative interaction of movements could be taken as a marker for neural fusion (Wang et al., 2024), it could also be the result of neural co-articulation of two separate plans initiated upstream.

Findings in humans and non-human primates stand in sharp contrast to the strong evidence from rodents, which aligns closely with the notion of neural fusion. Here neural activity in the striatum have been shown to fully reflect the intricate kinematics of a highly overtrained habitual sequence and was disrupted by lesioning of the dorsolateral striatum (Kawai et al., 2015; Dhawale et al., 2021; Thompson et al., 2024). A key difference from studies in non-human primates and humans is that rodents are typically trained on sequences in a highly rigid manner: movement elements are repeated habitually, without being recombined in different orders or performed with varying timings. When variability is introduced – such as requiring flexible use of the same elements alongside an overtrained sequence – the control of the sequence shifts from the striatum to the cortex (Mizes et al., 2024). This capacity for trial-by-trial flexibility is central to most human studies and to intricate, smoothly executed behaviours such as handwriting and typing, and may explain why neural fusion does not occur for most skilled sequential behaviours in primates despite the outward behavioural fusion of movement trajectories.

One possibility is that different neural representations of individual movements that we observe during planning shift towards integrated representations during execution. Neuroimaging findings revealed that during planning, premotor and parietal regions have been shown to encode individual features of a sequence such as movement order and timing independently whilst during execution, these regions dynamically shift to integrate these features into a joint sequence representation (Yewbrey et al., 2023). Therefore, individual sequence elements may be maintained during preparation but integrated during execution to produce smooth, fused action. In the current study, it could mean that during planning, individual movements are encoded separately, but as execution unfolds, specifically in highly fused sequences, these elements are combined into a higher-level joint representation. Fusion, then, may reflect a hierarchical shift from high-level modular to integrated control trial-by-trial.

Although we manipulated fusion in our study, complete average kinematic fusion between adjacent sequence elements (Fusion index equal to or close to 1.0) was only observed in a small subset of participants (Figure 5). Various studies have used multi-day experiments to achieve fusion (Sosnik et al., 2004; Sporn et al., 2022) to increase repetition and boost learning through sleep consolidation (Korman et al., 2003). Our study builds on Sporn et al. (2022), who showed that movement fusion increased over two days of practice before plateauing on the third. We adapted their sequential task but modified the design by using smaller, more closely spaced targets, more similar to handwriting. In the right- and obtuse-angle groups, we observed slightly higher degrees of fusion than previously reported for sequential reaching movements (Sosnik et al., 2004; Sporn et al., 2022). Note that the latter computed the fusion index cumulatively across seven movements. Whilst smaller amplitude movements may have facilitated movement fusion due to the spatial proximity of the sequence elements, there is a possibility that only consistent and complete behavioural fusion of the adjacent elements leads to neural fusion into a new motor representation. Importantly, natural sequential skills such as handwriting are seldom fully kinematically fused across all elements, making it unlikely that they consolidate into entirely distinct movement primitives. The current task is representative of such behaviours, as it captures the partial but not complete fusion characteristics of natural sequential behaviours.

While some degree of fusion was present at the outset of the study across all groups, fusion scores became less variable over time only for the right-angle group which showed the most consistent fusion across participants and participants exhibiting higher fusion in the median split across the three groups. This suggests that the change in fusion observed in these two groups reflects a learning-related effect, rather than simply representing biomechanical coarticulation that is present from the outset. Sosnik (2007) suggests that coarticulation is a slow process that occurs throughout training as a new motion planning strategy is acquired, possibly reflecting a change in the internal representations of the sequence. However, in our study the pre-ordering of the discrete constituent elements in the sequence was most pronounced in the right- and obtuse-angle groups, where the most relative fusion occurred. This suggests that the optimisation might not be achieved through a new motor sequence planning strategy per se, but the optimisation of already existing planning strategies, e.g. the consolidation of a CQ gradient.

Studies have observed significant individual variability in fusion. For example, Sosnik (2004) provided additional training sessions for participants who did not merge their movements to see whether this would induce fusion, yet two out of nine participants still performed the movements separately despite the extensive training. This suggests that while introducing task constraints, such as manipulating angle (Sosnik et al., 2004, 2007), manipulating reward (Sporn et al., 2022) or increasing accuracy demands (e.g., (Fitts’, 1954; Sosnik et al., 2007), can influence fusion to some extent, some individuals may not fuse their movements, and not be susceptible to a training intervention.

A key feature of skilled sequence production is the ability to initiate movements fluently and perform them accurately. Our present findings indicate that CQ is linked to more skilled performance more generally. Previous research in typing movements has shown that a larger position-dependent difference between CQ patterns influences subsequent performance (Kornysheva et al., 2019; Mantziara et al., 2021). The regression analyses with performance measures as predictors revealed that participants with a faster sequence initiation time and higher accuracy had a more pronounced gradient (ProbeRT), an effect also previously found in discrete typing-like sequences (Mantziara et al., 2021). Similarly, a higher fusion score was linked to a more pronounced CQ gradient (ProbeError), however this relationship was not upheld in the correlational results. Contrary to our hypothesis, we did not find lower fusion to be associated with a more pronounced CQ. The acute group had a diminished CQ gradient, along with a lower sequence accuracy during the test phase compared to the right-angle and obtuse groups. Moreover, the acute angle may have led participants to prioritise precision over fluidity, hindering skilled sequence performance. Overall, these findings suggests that a larger position-dependent gradient between sequence elements during planning leads to better performance in skilled sequences with smooth and continuous kinematics.

We note that in tasks such as target-jump paradigms, where the target jumps while the movement is underway, participants plan a new movement from the original target to the new target (Flash and Henis, 1991; Kashefi et al., 2024) and by vector summation they will arrive at the correct location. In general, they will execute the second sub-movement before the first has terminated, i.e., it is a form of fusion. The fusion described in this experiment is more complex, because it is not possible to simply vectorially sum unfused movements and still pass through all the targets. Rather, the participants are required to change the trajectories of the constituent sub-movements to successfully perform the task, a process which likely takes time, and varies across participants.

In conclusion, handwriting-like movements are organised in advance before the sequence is executed. The CQ of upcoming sequence elements was stronger in participants who performed continuous sequences from memory with higher skill – demonstrated by faster initiation, greater accuracy and larger kinematic fusion. This is consistent with previous findings in discrete sequences such as typing (Kornysheva et al., 2019; Mantziara et al., 2021) and indicates that the automatic pre-ordering of upcoming movements is a beneficial feature of general motor skill.

## Author contributions

H.W., J.F., J.G. and K.K. designed research; H.W. performed research; H.W., J.F. and K.K. analysed the data; H.W. and K.K. wrote first paper draft; H.W., J.F., J.G. and K.K. edited the paper draft.

## Acknowledgements

With thank the undergraduate students Christopher Chong, Amelia Carter, Holly Clark, Charlotte Lemon, Archie Robertson, Florence Sargent, Mohammed Shakeel for helping with part of the data collection, Brandon Ingram and Dagmar Fraser for technical assistance with the tablet and Coen Zandvoort for comments on earlier versions of the manuscript. This work was supported by Academy of Medical Sciences Springboard Award (SBF006/1052 to K.K.) and the UKRI Future Leaders Fellowship (MR/Y016467/1 to K.K).

